# Adjuvant conditioning enhances neutrophil function while inducing a suppressive peritoneal macrophage phenotype

**DOI:** 10.1101/2025.04.15.648954

**Authors:** Thais Boccia, Victor Fattori, Matheus Deroco Veloso da Silva, Nathan L. Asquith, Weikang Pan, Michael S. Rogers, Ivan Zanoni, Alex G. Cuenca

**Author notes:** Corresponding Author: Alex G. Cuenca, MD PhD, 300 Longwood Ave, Fegan 3, Department of Surgery, Boston Children’s Hospital, Boston, MA 02115.

## Abstract

Adjuvants are widely used to boost the immune response during vaccination protocols. Our group has previously reported that repeated intraperitoneal administration of alum in mice, known as adjuvant conditioning (AC), creates an immunosuppressive environment that delays allogeneic graft rejection through NLRP3-dependent MDSC expansion. However, little is known about the effects of AC on the reprogramming of peritoneal cavity cells, particularly the different peritoneal macrophage populations and the effects in the adaptive immune response. We found a population-specific immune response to alum, with small peritoneal macrophages (SPMs) being more prone to inflammasome activation than large peritoneal macrophages (LPMs) *in vitro*. *In vivo*, alum exposure led to NLRP3-dependent macrophage disappearance reaction (MDR) of LPMs, which could be explained by aggregate formation and migration to the omentum. AC also induced the reprogramming of resident macrophages and infiltrating monocytes towards a less inflammatory state, making them more vulnerable to bacterial infections, but recruited neutrophils with enhanced killing ability. This suggests that AC may influence both innate and adaptive immunity in distinct ways, reprogramming cells to different profiles, indicating its potential as an immunosuppressive treatment for autoimmune diseases and transplant rejection.

## Introduction

Transplant medication regimens attempt to achieve a delicate balance between immunoregulation of recipient alloimmunity and immunity to infection. However, the currently employed medications have deleterious effects on both adaptive and innate immunity, including unintended effects on the depletion or inhibition of immunoregulatory cell populations that can suppress alloimmunity (1–3). Therefore, therapeutics designed to immunomodulate the adaptive immune responses to prevent transplant-associated rejection but retain immunity to infection are needed.

Adjuvants are crucial components of vaccine preparations and are designed to enhance the immune response to targeted antigens (4). Many adjuvants, such as alum or monophosphoryl lipid A, can directly engage toll-like receptors (TLRs), thereby promoting the robust activation of dendritic cells and enhancing antigen presentation to T cells (5–8). Immunization via the intraperitoneal (i.p.) route is valuable for studying immune responses in animals due to the unique microenvironment of the peritoneal cavity, which is rich in immune cells, particularly peritoneal macrophages. Although most solid organ transplants are intra-abdominal, there is very little data on the role of peritoneal immunity in abdominal organ transplants.

In mice, peritoneal macrophages are comprised of distinct populations that exhibit unique characteristics and functions. They are primarily categorized into resident and recruited macrophages (9). Resident macrophages, maintain a stable presence in the peritoneal cavity and are crucial for tissue homeostasis and initial immune responses (10–12). In contrast, recruited macrophages arise from monocytes in response to inflammatory signals and play key roles in acute immune reactions, particularly during infection or injury (9, 11, 13). Within the resident population of peritoneal macrophages, two distinct subpopulations can be identified among the CD11b+ cells: the large peritoneal macrophage (LPM), characterized by high levels of F4/80, a classical macrophage marker, and low levels of MHC II, and the small peritoneal macrophage (SPM), which exhibits low F4/80 and high MHC II expression (9, 14–18). These subpopulations have been extensively studied, revealing specific cell surface markers, transcription factors, and their origins, as well as their roles during inflammatory responses and bacterial infections. Importantly, they also contribute to the activation of the adaptive immune response.

Our group has shown that repeated i.p. administration of alum, termed adjuvant conditioning (AC), establishes an immunosuppressive environment that delays allogeneic graft rejection (19). This immunosuppression is mediated by the expansion of myeloid derived suppressor cells (MDSCs) dependent on NLRP3 activation and IL-1 signaling (19, 20). However, there are no reports of how AC can reprogram the cells of the peritoneal cavity, and if this initial recognition of alum by the peritoneal macrophages is involved in the effects of AC in the adaptive immune response. Here we demonstrate that resident peritoneal macrophages exhibit distinct responses to alum *in vitro*, with small peritoneal macrophages (SPMs) showing a greater propensity for inflammasome activation compared to large peritoneal macrophages (LPMs). *In vivo*, exposure to the adjuvant induced reprogramming of macrophages and monocytes towards a less inflammatory phenotype, resulting in heightened susceptibility to bacterial infection, but at the same time recruited neutrophils with higher killing ability. This suggests that the conditioning process affects both innate and adaptive immunity in different ways, which raise the potential for using this reprogramming approach as an immunosuppressive treatment for conditions such as autoimmunity and transplant rejection.

## Methods

### Mice

For the study, 8–10-week-old males C57BL/6J (H-2b), OT-II (B6.Cg-Tg(TcraTcrb)425Cbn/J), and NLRP3^-/-^ (B6.129S6-Nlrp3tm1Bhk/J) mice were purchased from The Jackson Laboratory. All mice were kept under specific pathogen free conditions at Boston Children’s Hospital and experimental protocols were approved by the Institutional Animal Care and Use Committees of Boston Children’s Hospital.

### Cell isolation and culture

Mice were euthanized by CO inhalation, and the abdominal cavity was aseptically exposed. The peritoneal cavity was washed with 10mL of ice-cold phosphate-buffered saline (PBS) with 10% of FBS. A sterile syringe with a 25-gauge needle was used to introduce the lavage fluid into the peritoneal cavity, and gentle massage was applied to ensure thorough washing of the peritoneal surfaces. The lavage fluid was then collected and transferred into a sterile 15 mL conical tube. The collected lavage fluid was immediately centrifuged 300 × g for 5 minutes at 4°C to pellet the peritoneal cells. The supernatant was discarded, and the cell pellet was resuspended in an appropriate volume of RPMI 1640 medium (ThermoFisher Scientific) containing 10% fetal bovine serum (Corning), 200μg/mL penicillin (ThermoFisher Scientific), 200 U/mL streptomycin (ThermoFisher Scientific), and 0.05mM 2-mecaptoethanol (Sigma-Aldrich) for downstream analyses (flow cytometry, cytokine assays, and immunofluorescence). The total cell count was determined using a hemocytometer, and cell viability was assessed by trypan blue exclusion.

To identify the cell populations in the omentum and examine the potential formation of aggregates following intraperitoneal (i.p.) alum injections, omental tissues were harvested at the time points indicated in the figures. The samples were immediately placed in a digestion solution containing DNase I (0.25 mg/mL) (Roche) and Liberase (0.05 mg/mL) (Roche) in DMEM low glucose (11054020, Thermo Fisher Scientific). The omentum was incubated with gentle agitation at 37°C for 30 minutes, while the aggregates were incubated for 5 minutes under the same conditions. A detailed protocol for isolating cells from both the omentum and aggregate structures is provided in Vega-Perez (2021) and Ferriz (2023) (21, 22). Neutrophils were isolated from either alum-treated or thioglycolate-elicited C57BL/6 mice (23, 24) and purified as described elsewhere (24).

### Cell viability

Cell viability was assessed using a dual-staining method with ethidium bromide (EtBr) and acridine orange (AO), which allows for the discrimination between viable and necrotic cells (25). Following the incubation of peritoneal macrophages with LPS (50ng/mL) and alum Imject (Thermo Fischer Scientific) (concentrations indicated in the figures), cells were washed twice with PBS and resuspended in a mixture of EtBr (Sigma Aldrich) and AO (Sigma Aldrich) was prepared at final concentrations of 50μg/mL for both EtBr and AO. Cells were then incubated at room temperature for 5 minutes in the dark to allow for adequate dye uptake. After incubation fluorescence microscopy was performed using a EVOS M7000 Imaging System (Thermo Fischer Scientific) equipped with appropriate filters for detecting ethidium bromide and acridine orange fluorescence. Viable cells appeared with a uniform green fluorescence in the cytoplasm and nucleus when observed under the green fluorescence channel (excitation: 488 nm, emission: 530 nm). Necrotic cells were stained with red fluorescence, indicating the presence of EtBr, which is impermeable to live cells but stains cells with compromised membranes. Cell counts were performed for each category viable and necrotic, and results were expressed as a percentage of the total cell population. Additionally, cell viability under the same conditions was measured using the CyQUANT™ LDH Cytotoxicity Assay (Thermo Fischer Scientific) following the manufacturer’s instructions.

### Drug administration

For adjuvant administration, mice received either saline or alum Imject (8mg in 200 μL) (Thermo Fischer Scientific) intraperitoneally every other day for three times. For IL-1 signaling blockade, KINERET® (Anakinra) was injected intraperitoneally at a dose of 30mg/kg at time points described in each figure.

### CD4 T cell proliferation assay

To assess CD4 T cell proliferation, naïve CD4+ T cells were isolated from OT-II mouse spleens using EasySep mouse naïve CD4+ T cell isolation kit (StemCell) and labeled with CellTrace CFSE (ThermoFisher Scientific). Macrophages and/or monocytes were isolated from alum-treated or saline-treated animals using an BD FACSAria™ II Cell Sorter and cultured with naïve CD4 T cells in the presence of 1μg/mL of OVAp 323-339 (InvivoGen) for 72h. Samples were collected on LSR II with BD FACSDiva v8.0.2 software (BD Biosciences). FlowJo v10 was used for flow data analysis.

### Flow cytometry

Single-cell suspensions were prepared from mouse peritoneal cavity, omentum or aggregates. The following antibodies were used for surface staining: CD11b (clone: M1/70; Invitrogen), Ly6C (clone: HK1.4; BioLegend), Ly6G (clone: 1A8; BioLegend), F4/80 (clone: BM8; BioLegend), MHC II (clone: M5/114.15.2; BioLegend), CD19 (clone: 1D3; BioLegend), CD11c (clone: N418; BioLegend), CD3 (clone: 17A2; Invitrogen), CD4 (clone: GK1.5; Invitrogen). Samples were fixed with eBioscience™ IC Fixation Buffer. Cell viability was determined Fixable Viability Dye eFluor™ 780 (eBioscience™), Zombie Violet™ or Zombie Aqua™ Fixable Viability (Biolegend). Samples were collected on LSR II with BD FACSDiva v8.0.2 software (BD Biosciences). FlowJo v10 was used for flow data analysis.

### Caspase-1 detection

Mice were injected with either saline or alum (8mg in 200uL) and euthanized by CO inhalation after 30 minutes. The lavage fluid was then collected and transferred into a sterile 15 mL conical tube. The collected lavage fluid was immediately centrifuged 300 × g for 5 minutes at 4°C to pellet the peritoneal cells. The supernatant was discarded, and the cell pellet was resuspended in an appropriate volume of RIPA lysis buffer (Thermo Fischer Scientific) containing protease and phosphatase inhibitors (Thermo Fischer Scientific). Total protein was assessed by BCA assay (Thermo Fischer Scientific). 20μg of total protein was mixed with loading buffer, boiled for 5 minutes at 90°C, and cooled to room temperature. SDS-PAGE gel electrophoresis was performed in NuPAGE 12% acrylamide gels submerged in tris buffer pH 7.4. Proteins were separated at 70V for 10 minutes, followed by 60 mins at 120V. The gel was assembled into a western blot sandwich per manufacturers’ instructions (BioRad), and the MW ladder and samples were transferred to a 0.45um PVDF membrane using a TransBlot Turbo transfer machine (BioRad) at 25V for 7 mins. The resulting western blot was washed in tris buffer and blocked in 3% BSA for 1 hour, followed by overnight incubation with primary rabbit anti-mouse caspase-1 antibody (#89322 Cell Signaling Technology). The blot was washed with tris and incubated with secondary rabbit anti-mouse-HRP (#7076S, Cell Signaling Technology) for 2 hours at room temperature. The blot was washed in tris and ECL cocktail (BioRad), applied per the manufacturers’ instructions, and exposed cumulatively for a maximum of five minutes on a Biorad ChemiDoc. Images were processed using ImageLab (BioRad).

### Immunofluorescence imaging

Total or sorted peritoneal macrophages/monocytes were plated overnight for adhesion onto cell imaging chambers (Lab-Tek #155383) at 37° C and 5% CO_2_. The supernatant was removed, and the adhered cells were treated as indicated in each figure. After treatment, cells were fixed with 4 % PFA for 30 minutes at room temperature, then blocked using 3% BSA for 2h. Cells were stained using primary antibodies for F4/80 AF647 (clone: BM8, BioLegend), purified rat anti-mouse MHC II (clone: M5/114.15.2, BioLegend), secondary antibody anti-rat AF594 (Thermo Fischer Scientific), and DAPI (Thermo Fischer Scientific). Following secondary antibody staining, cells were washed with PBS and incubated in PBS at 4 ° C until imaging. All fixed-cell microscopy was performed with a Zeiss AxioObserver laser scanning confocal microscope 880.

### Cytokine detection

Cytokine release in the supernatant was assessed using LEGENDplex™ Mouse Inflammation Panel 1 (740446; BioLegend). IL-β and IL-α concentrations were measured by Mouse IL-1 beta/IL-1F2 DuoSet ELISA (DY40105; R&D Systems™) and Mouse IL-1 alpha/IL-1F1 Quantikine ELISA Kit (MLA00; R&D Systems™).

### ROS and Nitric oxide detection

Nitric oxide production by macrophages cultured *in vitro* was measured by Griess Reagent Kit (Invitrogen). Reactive oxygen production by macrophages cultured *in vitro* was measured by ROS-Glo™ H_2_O_2_ Assay (Promega).

### Antigen processing

Antigen processing by the macrophages and monocytes was assessed by culturing the harvested cells in a 4μg/mL solution with DQ™ Ovalbumin (Invitrogen) for 4h. The antigen processing levels were analyzed either by flow cytometry or by fluorescence microscopy.

### Phagocytosis

Phagocytic activity was assessed by incubating cells with Zymosan A (*S. cerevisiae*) BioParticles™, Alexa Fluor™ 488 conjugate (Thermo Fischer Scientific) or Zymosan A from *Saccharomyces cerevisiae* (Sigma Aldrich). Zymosan, a commonly used particle for assessing phagocytosis, was opsonized using mouse serum, and added to cell cultures for 30 minutes to evaluate the uptake by phagocytic cells. The cells were washed with ice-cold PBS and then analyzed either by flow cytometry or by fluorescence microscopy.

### Bacterial infection – *E. coli*

*E. coli* strain (*Escherichia coli* GFP – ATCC 25922GFP) was grown overnight in Tryptic Soy Broth (Sigma Aldrich) at 37°C with shaking (250 rpm). The bacteria were cultured to exponential growth (OD600 of 0.5) and then harvested by centrifugation at 4,000 × g for 10 minutes, washed twice with PBS, and resuspended in RPMI 1640 10% FBS. The bacterial concentration was determined by OD_600_ and colony-forming unit (CFU) assay to ensure accurate dosing.

Macrophages were infected with *E. coli* at an MOI of 5 by adding the bacterial suspension to the macrophage monolayer in RPMI 1640 10% FBS. The cells were centrifuged at 100 × g for 5 minutes and incubated at 37°C for 1 hour, allowing bacterial uptake. After the infection period, unbound bacteria were removed by washing the macrophages three times with pre-warmed PBS to reduce background contamination. Fresh culture medium without antibiotics was then added to the cells and macrophages were incubated for 24 hours at 37°C in a 5% CO incubator. At specific time points, macrophages were harvested for further analysis, including reactive oxygen specied (ROS) measurement, nitric oxide (NO) production, and bacterial load determination. At 24h post-infection, macrophages were collected for analysis. For bacterial load determination, cells were lysed in 0.1% Triton X-100 (Sigma Aldrich) in PBS, and serial dilutions were plated on LB agar plates to quantify intracellular bacteria. For cytokine analysis, supernatants were collected and stored at −80°C until analysis by ELISA or multiplex assays.

In additional experiments, macrophages and/or neutrophils were infected in a 15 mL conical tube in order to allow staining for surface markers using flow cytometry. Alternatively, cells were fixed with 4% paraformaldehyde for 30 minutes and stained with antibodies against macrophage markers. The uptake of bacteria was visualized and quantified by flow cytometry.

### Data Analysis and Statistics

Statistical analysis was performed using GraphPad Prism, version 10. A one-way analysis of variance was used for multiple comparisons. A p-value of <0.05 was considered a statistically significant difference.

### Data availability

Data acquired specifically for this study are available within the article itself and the supplementary materials. Experimental protocols and additional details regarding methods employed in this study will be made available upon reasonable request to the corresponding author.

## Results

### Resident peritoneal macrophages respond to alum stimulation *in vitro*

Previous studies investigating the effects of alum stimulation on peritoneal macrophages have primarily focused on cells recruited by thioglycolate (26–28). To assess the response of resident macrophages, we cultured steady-state peritoneal cavity resident cells followed by stimuli with both LPS and alum. We found that increasing concentrations of alum induced cell death in these macrophages, as indicated by increased ethidium bromide staining and LDH release (**Fig 1A-B and Supplemental Fig 1**). This effect was more pronounced in LPS-primed macrophages, which is consistent with previous reports on inflammasome activation (29, 30). Notably, resident macrophages were capable of producing IL-1α and IL-1β in response to alum, even without prior LPS priming (**Fig 1C-D**). Although IL-1β levels were low in the absence of LPS, this finding suggests a primed state for inflammasome activation in these cells.

**Figure 1:**
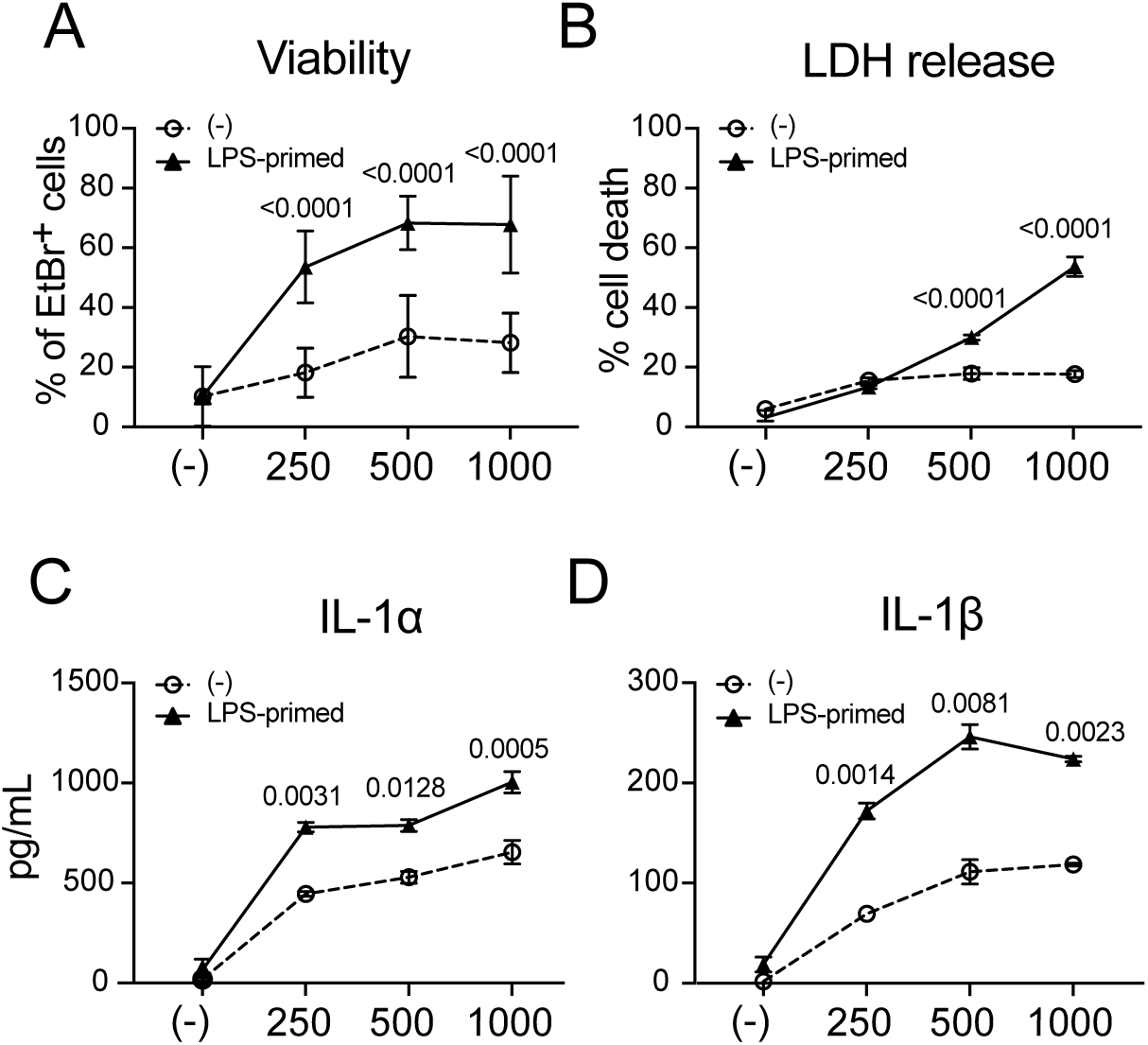
Resident peritoneal macrophages respond to alum stimulation *in vitro*. 3×10^5^ peritoneal cells from C57BL/6 mice were plated on 96-well plates overnight. After adhesion, non-adherent cells were washed out with cold PBS, and cells were primed with LPS (50ng/mL) for 2h and subsequently stimulated with alum (Imject) at 250, 500 or 1000μg/mL for 6h. Cells were then analyzed for viability using either a fluorescence microscope or LDH release. **A:** Percentage of cell death measured by ethidium bromide and acridine orange staining. **B:** LDH release in the supernatant **C:** IL-1α release in the supernatant **D:** IL-1β release in the supernatant. Data shown represent three or more experiments and are expressed as mean±SEM; Student’s t-test was used for analysis; P values are indicated in the graphs.

### Peritoneal macrophages are composed of two different populations

As reported by other groups, previous studies have identified multiple cell types, including B cells, dendritic cells, and macrophages, within the peritoneal cavity (9). Peritoneal macrophages can be further categorized into two populations based on their expression of surface markers: large peritoneal macrophages (LPMs) and small peritoneal macrophages (SPMs). LPMs are CD11b^+^F4/80^high^MHCII^low^, while SPMs are CD11b^+^F4/80^low^MHCII^high^ (9, 11, 14, 31). We observed a predominance of LPMs over SPMs in the peritoneal cavity of C57BL/6 mice, as confirmed by immunofluorescence (**Fig 2A-B**) and flow cytometry (**Fig 2C and Supplemental Fig 2A**). To investigate potential differences in the responses of these two macrophage populations to alum, we sorted and stimulated the macrophages with LPS and alum. Our findings revealed that SPMs are more susceptible to inflammasome activation and cell death by pyroptosis compared to LPMs. This was evident from increased ethidium bromide staining (**Fig 2E-F**) and LDH release in the presence of alum (**Fig 2E-F**). Additionally, SPMs produced higher levels of IL-1α and IL-1β than LPMs (**Fig 2G**), suggesting distinct inflammatory programming between these two macrophage subsets.

**Figure 2:**
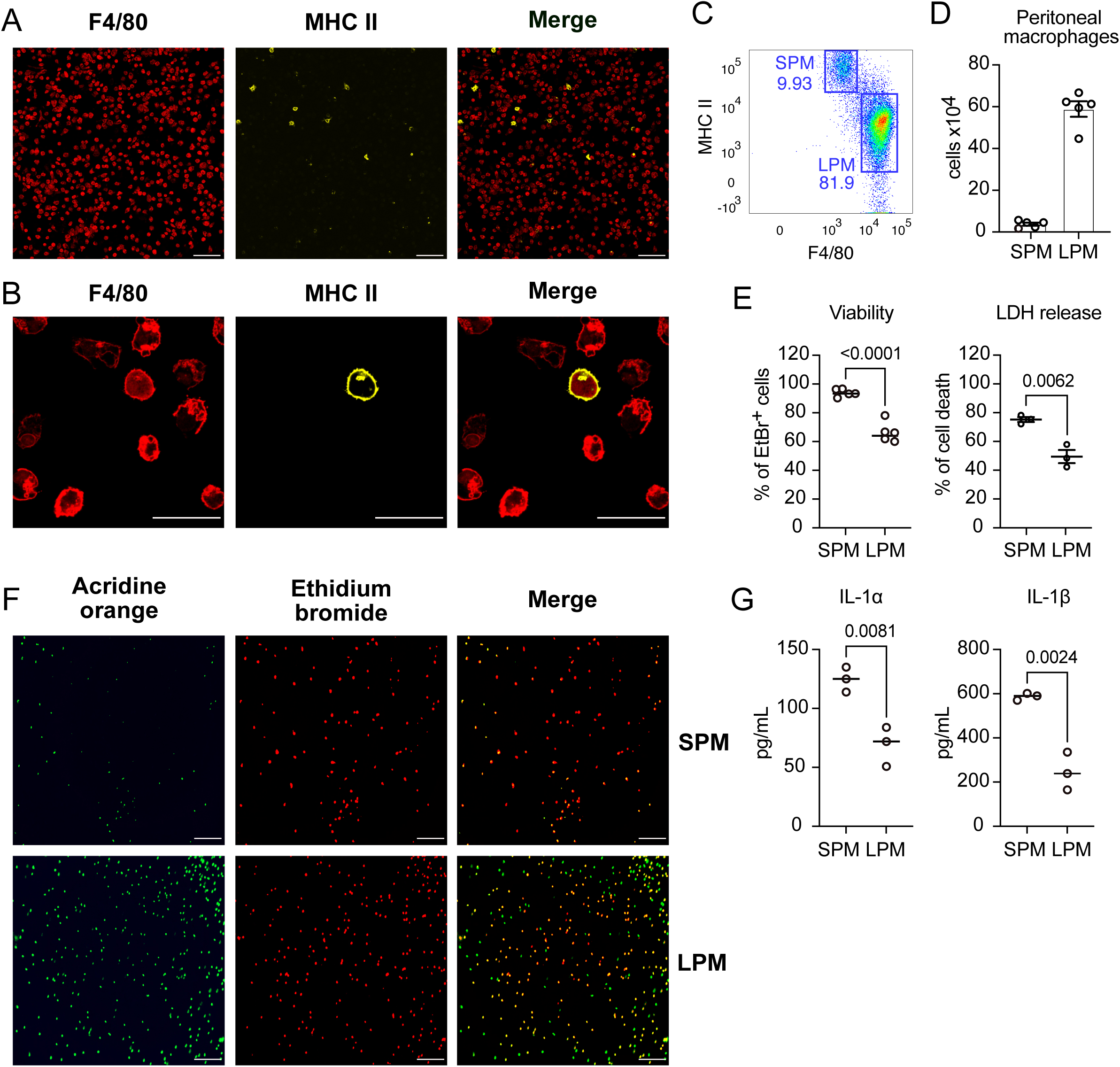
Peritoneal macrophages are composed of two different populations. **A:** 5×10^5^ peritoneal cells from C57BL/6 mice were plated on to imaging chambers overnight. After adhesion, non-adherent cells were washed out with cold PBS, and cells were stained with antibodies for F4/80 and MHC II for immunofluorescence microscopy. Scale bars represent 10μm. **B:** Magnified view of the cell population depicted in A. Scale bars represent 10μm. **C:** Flow cytometry plots of CD11b^+^Ly6G^neg^ macrophage populations in a steady state peritoneal cavity, highlighting SPMs (small peritoneal macrophage F4/80^low^MHC II^high^), and LPMs (large peritoneal macrophage F4/80^high^MHC II^low^). See Supplemental Figure 1A for the full gating strategy. **D:** Total count of SPMs and LPMs in a steady state peritoneal cavity. **E:** Sorted SPMs and LPMs plated at 2×10^5^ cells/well, were primed with LPS (50ng/mL) for 2h and subsequently stimulated with alum (Imject) at 500μg/mL for 6h. Cells were assessed for viability and the percentage of EtBr^+^ cells and LDH release in the supernatant are depicted. **F:** IL-1α and IL-1β release in the supernatant of sorted SPMs and LPMs treated with LPS, and alum detected by ELISA. Data shown represent three or more experiments and are expressed as mean±SEM; Student t-test was used for analysis; P values are indicated in the graphs.

### AC alters cellular composition of peritoneal innate immunity

Our research group has previously shown that repeated intraperitoneal injections of alum, a process called adjuvant conditioning, can expand MDSCs (19, 20). These cells influence the adaptive immune response by limiting the proliferation of T cells and promoting immunotolerance, which delay allograft rejection (20). Because peritoneal macrophages are the first cells to respond to alum after injection (32), we hypothesized that AC might affect these cells. To test this hypothesis, we administered alum intraperitoneally three times every other day (**Fig 3A**). We then examined various immune cell populations within the peritoneal cavity 24h after the last injection.

**Figure 3:**
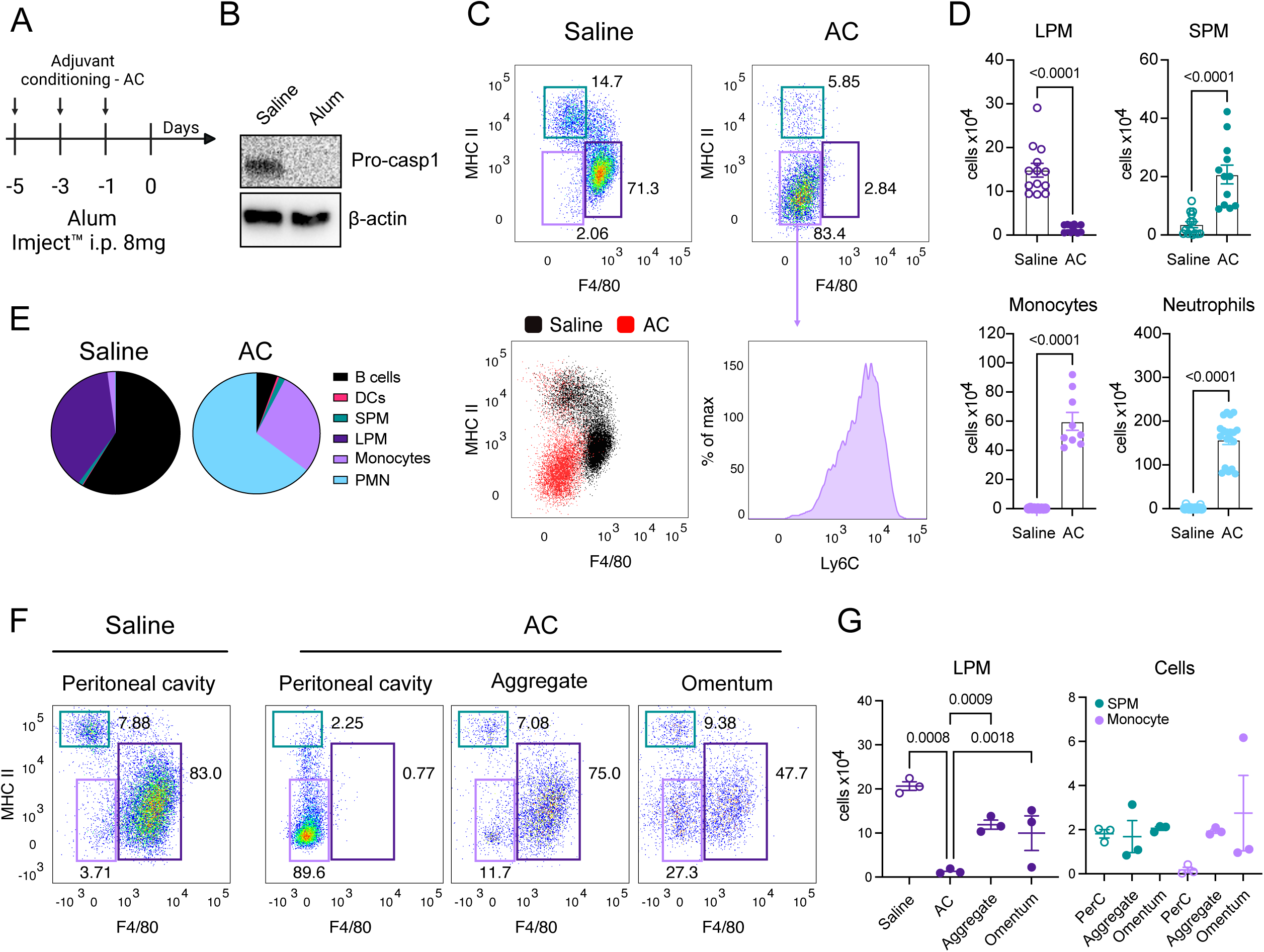
Adjuvant conditioning (AC) changes the populations of immune cells in the peritoneal cavity *in vivo*. **A:** Experimental design for AC. 8-10-week-old male C57BL/6 mice were injected intraperitoneally (i.p.) with either alum (Imject) (8mg in 200µL) or saline three times, every other day. **B:** Western blot analysis of pro-caspase-1 processing by peritoneal cells 30 minutes after alum or saline injection i.p. **C:** 24 hours after the last alum or saline injection, the peritoneal lavage was collected and phenotyped by flow cytometry. SPMs and LPMs are shown in saline or alum-injected (AC) mice. In the bottom left plot, both saline and AC macrophage plots are overlaid to highlight the differences in the populations, and the bottom right plot represents Ly6C expression in the infiltrating CD11b^+^F4/80^low^MHC II^low^ population of monocytes. **D:** Total cell number of LPMs, SPMs, monocytes and neutrophils in the peritoneal cavities of either saline or AC-treated mice. **E:** Fraction of all populations phenotyped in the peritoneal cavities of either saline or AC-treated mice. **F:** Comparison of macrophages and monocyte populations identified in the peritoneal cavity and also found in aggregates and omentum of either saline or AC-treated mice. **G:** Total cell number of LPMs, SPMs, monocytes and neutrophils in the peritoneal cavities, aggregates and omentum of either saline or AC-treated mice. Data shown represent three or more experiments and are expressed as mean±SEM; Student t-test was used for analysis; P values are indicated in the graphs.

Our results revealed that AC led to caspase-1 activation as early as 30 min after i.p. injection, as shown by the loss of pro-caspase-1(**Fig 3B**). In addition, alum induced MDR as observed by the reduction of LPMs as well as increased SPMs (**Fig 3C-D**) (33, 34). Interestingly, a mixed population of Ly6C^+^ and Ly6C^-^ monocytes (CD11b^+^F4/80^low^MHCII^low^) migrated into the peritoneal cavity, indicating that alum induces the recruitment of a novel and distinct monocyte population (**Fig 3C-D**). Overall, AC caused a major change in the immune cell landscape (Fig 3E). Specifically, we observed a dramatic reduction in B-cells, LPMs, and DCs and a subsequent increase in neutrophils and monocytes (**Fig 3E**) (32, 34, 35). Several studies have reported that resident peritoneal macrophages migrate to various organs, such as the omentum (18), or form aggregate scaffolds within the inner peritoneal layer following an inflammatory stimulus (18, 21, 36). We then wanted to investigate whether AC could also induce a similar phenomenon. Here, we report for the first time that AC not only promotes the formation of resident macrophage aggregates but also stimulates these cells to migrate to the omentum, contributing to the observed MDR (**Fig 3F-G** and **Supplemental Fig 3**).

### AC alters peritoneal cavity immune cell landscape

Given that the effects of AC primarily involved adaptive immune cells, particularly T cells (20), and resulted in delayed allograft rejection (19, 20), we aimed to investigate the peritoneal populations 7 and 14 days following the final injection. Our findings indicate that, 14 days after the conditioning protocol, LPMs return to the peritoneal cavity, likely due to the proliferation of the residual population or resolution of inflammation (**Fig. 4A-C and Supplemental Fig 3**). The infiltrating monocytes peak 24 hours after the last injection but return to baseline levels by day 14. During this period, they likely differentiate into SPMs, which may explain the variation in total SPM numbers observed between day 1 and day 14. Neutrophils also peak on day 1 but unlike monocytes, return to near homeostatic levels by day 7 (**Fig. 4A-C**). Also, no macrophage aggregates were found 7 days and 14 days after the conditioning protocol, suggesting that the peritoneal cavity partially to a resolved inflammation state as reported in studies using different inflammatory stimuli (14, 17). This indicates that AC changes the landscape of immune cells in the peritoneal cavity, and the resolution of the inflammatory insult takes place in up to 14 days.

**Figure 4:**
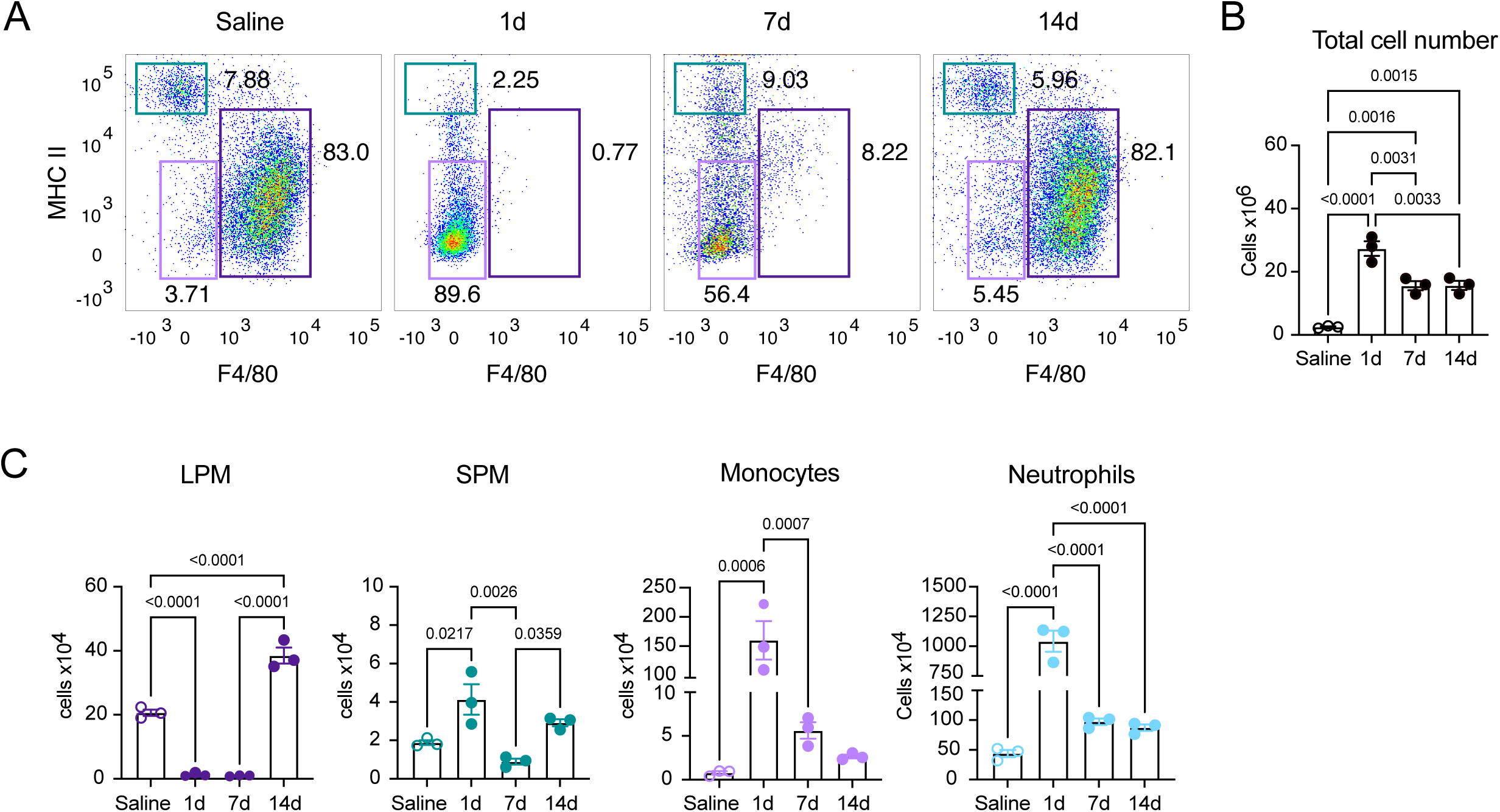
The effects of adjuvant conditioning (AC) on the peritoneal cell populations long term. **A:** Analysis of macrophages and monocyte populations in the peritoneal cavity of either saline or AC-treated mice at 1 day, 7 days and 14 days after the last injection. **B:** Total cell number in the peritoneal cavity of either saline or AC-treated mice at 1 day, 7 days and 14 days after the last injection. **C:** Total number of LPMs, SPMs, monocytes, and neutrophils in the peritoneal cavity of either saline or AC-treated mice at 1 day, 7 days and 14 days after the last injection. See supplemental Figure 4 for complete data on aggregates and omentum in all timepoints. Data shown represent three or more experiments and are expressed as mean±SEM; Student t-test was used for analysis; P values are indicated in the graphs.

### AC-induced macrophages and monocytes have a lower inflammatory profile

Having observed that AC dramatically changes the immune cell landscape in the peritoneal cavity, we next wanted to determine whether that would lead to any functional changes. To then assess the functional changes of these newly migrated peritoneal cavity cells, we then performed a zymosan phagocytosis assay in vitro from sorted LPMs and SPMs from saline-injected mice and SPMs and monocytes from AC-treated mice. As shown in **Fig. 5A-C**, steady-state LPMs exhibited significantly higher phagocytic activity compared to SPMs under the same conditions. The phagocytic capacity of AC-induced SPMs and monocytes was similar to that of steady-state SPMs (**Fig. 5A-C**). When stimulated with LPS *in vitro*, steady-state LPMs produced higher levels of IL-6, TNF-α, NO, and ROS. In contrast, this inflammatory profile was not observed in either steady-state SPMs or AC-induced SPMs and monocytes (**Fig. 5D-E**). To determine whether the effects of AC also influenced LPMs that had aggregated or migrated to the omentum, we isolated macrophages from these structures and cultured them in the presence of LPS. These macrophages also produced lower levels of inflammatory cytokines, NO, and ROS (**Fig. 5D-F**). Altogether, this indicates exposure to AC leads to a global immunosuppressive effect in macrophages that results in newly recruited and less inflamed immune cells in the peritoneal lavage after AC, as well as in the omentum.

**Figure 5:**
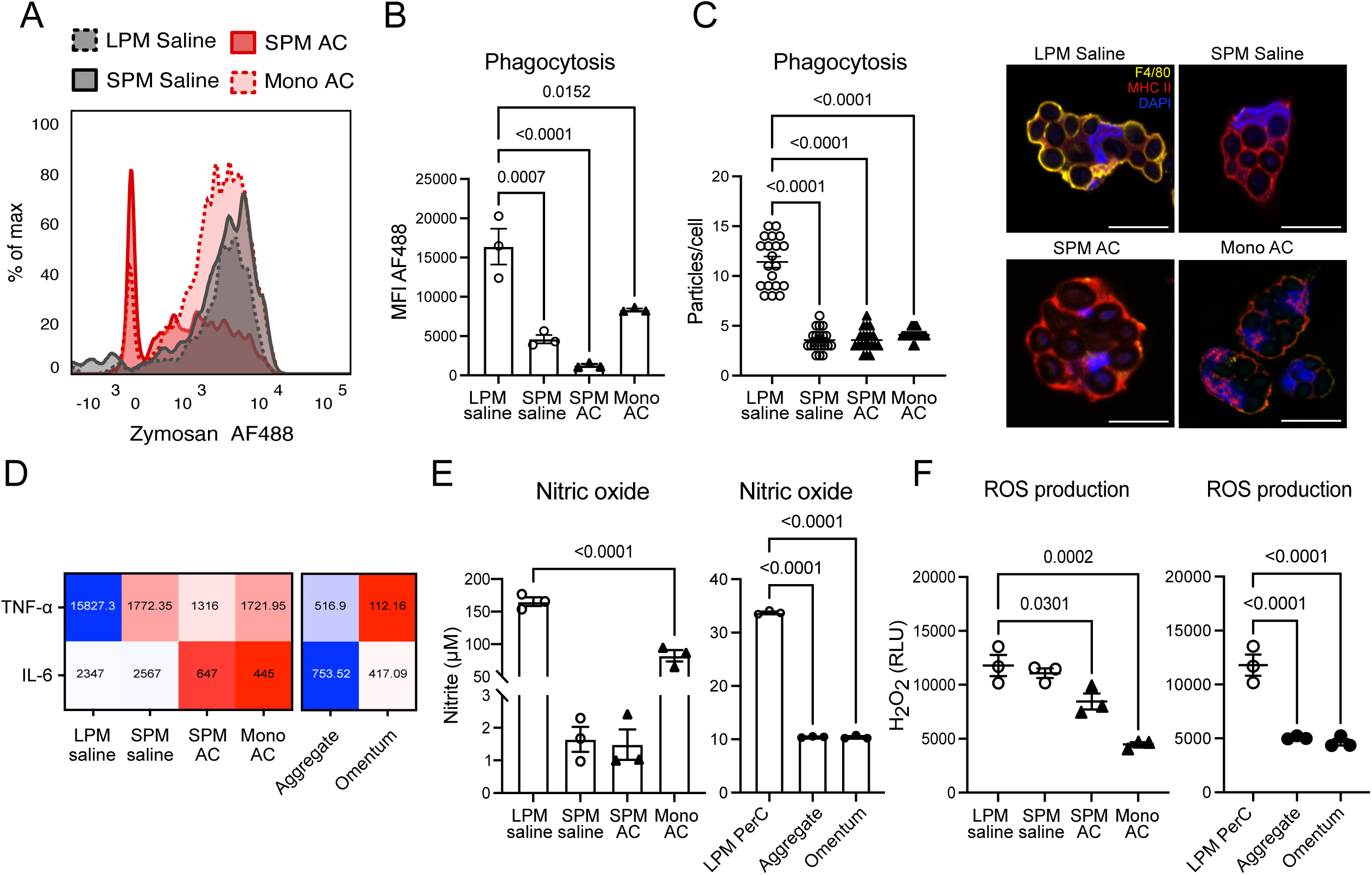
AC-induced macrophages and monocytes have a lower inflammatory profile. **A:** Histograms of AF488 conjugated Zymosan-A-treated LPMs, SPMs and monocytes from either saline or AC-treated mice. Cells were stained for flow cytometry phenotyping and analyzed for phagocytosis. **B:** Median of fluorescence intensity (MFI) of Zymosan- A-treated LPMs, SPMs and monocytes from either saline or AC-treated mice. **C:** Sorted SPMs, LPMs, and monocytes from saline or AC-treated mice were plated at 5×10^5^ cells/imaging chamber, and cultured with opsonized Zymosan-A for 30 min. Cells were then stained for immunofluorescence phenotyping and analyzed for phagocytosis. Scale bars represent 10μm. **D:** Sorted SPMs, LPMs, and monocytes from the peritoneal cavity (PerC), aggregates, and the omentum from saline or AC-treated mice were plated at 2×10^5^ cells/well of a 96-well plate and cultured overnight for adhesion. The supernatant was removed, and the cells were then stimulated with LPS (200ng/mL) for 24h. The supernatants were collected to measure TNF-α, IL-6, NO **(E),** and ROS **(F)**. Data shown represent three or more experiments and are expressed as mean±SEM; Student t-test was used for analysis; P values are indicated in the graphs.

### AC-induced SPMs and monocytes have poor antigen processing ability

Given that the AC-recruited cells exhibit less ability to induce an inflammatory response compared to steady-state resident macrophages, we turned to the adaptive immune response. To further assess the functionality of macrophages and monocytes induced by AC, we harvested total peritoneal cells from saline- or AC-treated mice and incubated them with DQ-OVA, which fluoresces upon proteolysis and is a marker for antigen presentation. Our results show that saline-treated macrophages have a greater ability to process OVA compared to AC-induced cells (**Fig 6A-B**). When examining each population individually, we found that steady-state LPMs are highly effective at processing antigens, outperforming both SPMs and AC-induced SPMs and monocytes (**Fig 6C-D**). However, despite their strong antigen-processing capability, LPMs were unable to activate OT-II CD4 T cells *in vitro*, likely due to their lack of MHC II expression. In contrast, both steady-state and AC-induced SPMs were able to activate T cells and promote proliferation (**Fig 6E**), suggesting that the presence of MHC II is more critical for CD4 T cell activation than the ability to process antigens.

**Figure 6:**
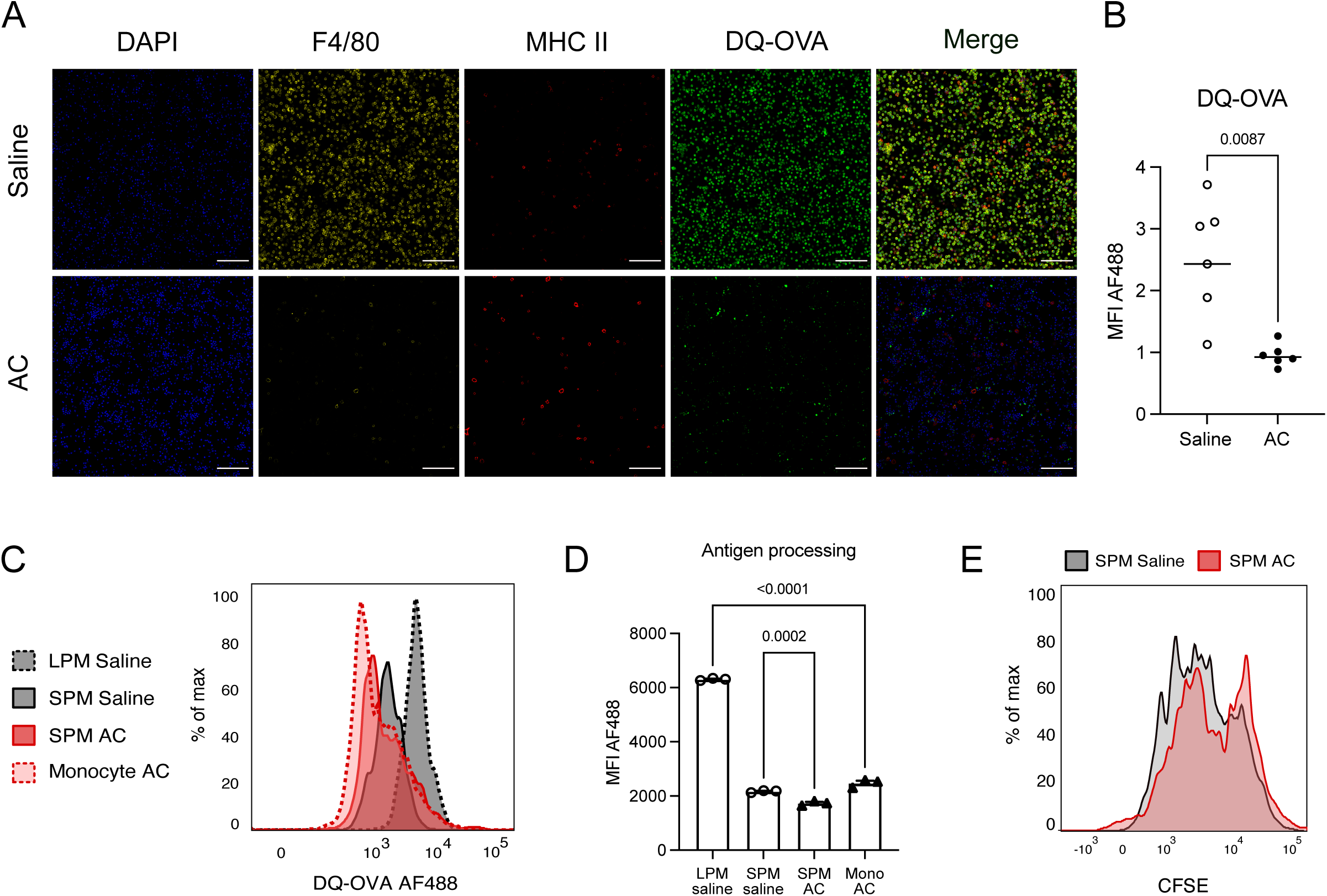
AC-induced SPMs and monocytes have poor antigen processing ability; however, SPMs still activate T cells. **A:** Total peritoneal cells from saline or AC-treated mice were plated at 5×10^5^ cells/imaging chamber and cultured with 4μg/mL DQ™ Ovalbumin for 4h. Cells were then stained for immunofluorescence phenotyping and analyzed for antigen processing. Scale bars represent 10μm. **B:** DQ™ Ovalbumin MFI of images taken from total peritoneal cells from saline or AC-treated mice. **C:** Histograms of AF488 DQ™ Ovalbumin-A-treated LPMs, SPMs and monocytes from either saline or AC-treated mice. **D:** DQ™ Ovalbumin MFI in LPMs, SPMs and monocytes from either saline or AC-treated mice. **E:** Histograms of CD4^+^Vα^+^ cells in a proliferation assay using isolated CD4^+^cells from spleens of OT-II mice cultured with OVAp (1µg/mL) and sorted SPMs from either saline or AC-treated mice. Data shown represent three or more experiments and are expressed as mean±SEM; Student t-test was used for analysis; P values are indicated in the graphs.

### NLRP3 activation is required for the distinct responses of LPMs and SPMs to alum

It is well established that stimulation of macrophages with alum activates the NLRP3 inflammasome, leading to the production of IL-1β (**Fig. 1D**) (29, 37). However, little is known about how the different populations of peritoneal macrophages respond to inflammasome activators. To investigate whether the differential production of IL-1α and IL-1β, as well as pyroptotic cell death in steady-state LPMs and SPMs involves NLRP3, we harvested total peritoneal cells from C57BL/6 and NLRP3-deficient mice. These cells were sorted into LPMs and SPMs and treated with LPS and alum *in vitro*. Our results showed that NLRP3^-/-^ LPMs and SPMs exhibited no differences in IL-1α production and LDH release (**Fig 7A-B**), suggesting that the observed differences in their responses to alum may be attributed to the differential expression of this inflammasome component. As anticipated, no IL-1β was detected in the supernatants of either NLRP3^-/-^ LPMs or SPMs.

**Figure 7:**
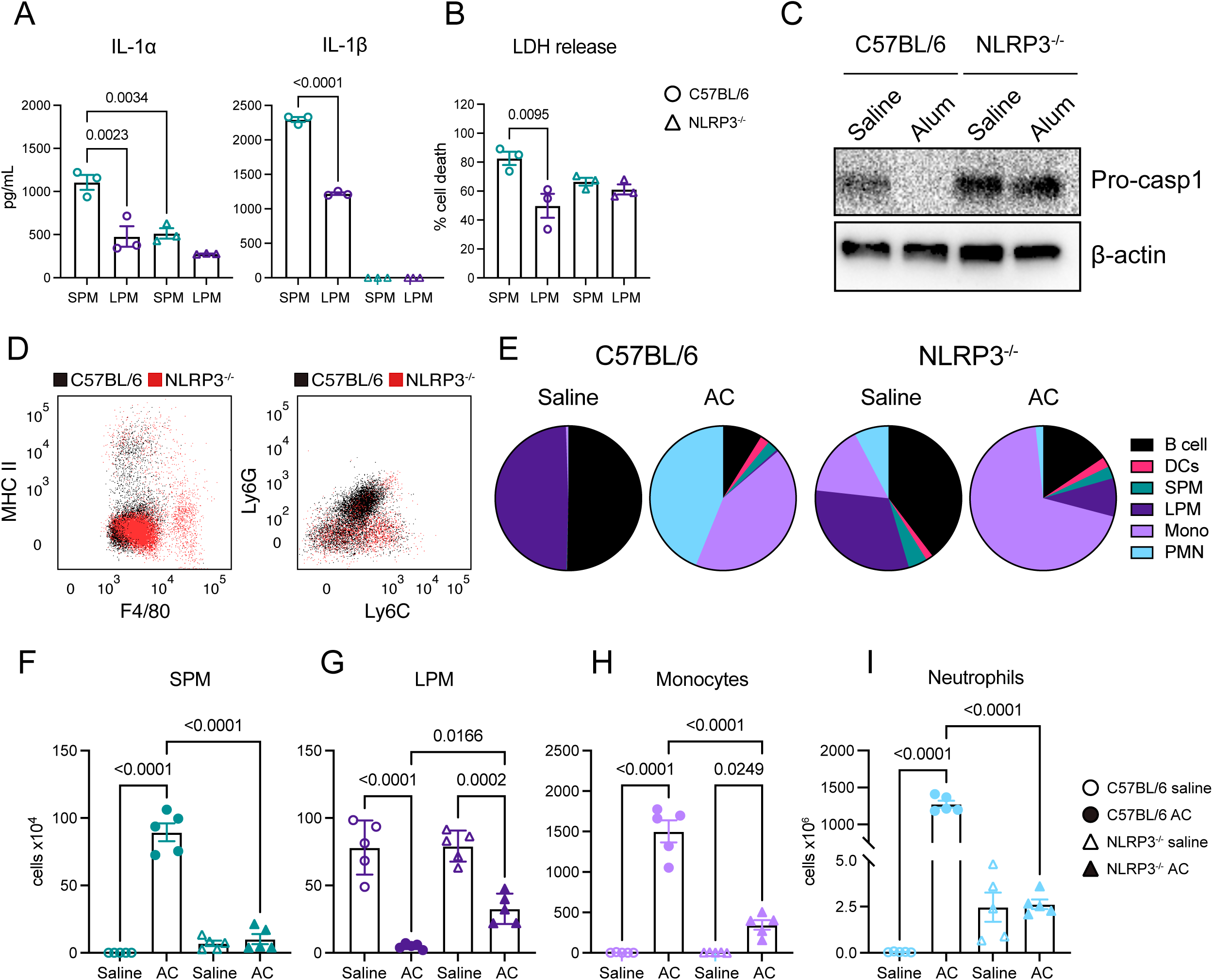
NLRP3 but not IL-1 signaling, is involved in AC-induced changes to the macrophage/monocyte populations in the peritoneal cavity *in vivo*. **A:** 3×10^5^ peritoneal cells from C57BL/6 and NLRP3^-/-^ mice were plated on to 96-well plates overnight. Cells were primed with LPS (50ng/mL) for 2h and subsequently stimulated with alum (Imject) at 500μg/mL for 6h. IL-1α and IL-1β release in the supernatant of peritoneal macrophages treated with LPS and alum. **B:** LDH release in the supernatant of peritoneal macrophages treated with LPS and alum in increasing concentrations. **C:** Analysis of pro-caspase-1 processing by peritoneal cells from C57BL/6 and NLRP3^-/-^ mice 30 minutes after alum or saline injection i.p. **D:** 24 hours after the last alum or saline injection, the peritoneal lavage of C57BL/6 and NLRP3^-/-^ mice was collected and phenotyped by flow cytometry. The plot to the left shows C57BL/6 and NLRP3^-/-^ mice AC macrophage plots overlaid to highlight the differences in the populations, and the right plot represents the infiltrating CD11b+Ly6G cells after AC in C57BL/6 and NLRP3^-/-^ mice. **E:** Fraction of all populations phenotyped in the peritoneal cavities of either AC- treated C57BL/6 and NLRP3^-/-^ mice. **F:** Total cell number of SPMs, LPMs **(G)**, monocytes **(H)** and neutrophils **(I)** in the peritoneal cavities of either either saline or AC- treated C57BL/6 and NLRP3^-/-^ mice. **J:** Total cell number of SPMs, LPMs, monocytes and neutrophils in the peritoneal cavities of either saline, AC, or AC+AK-treated C57BL/6 mice (AK: Anakinra 30mg/kg i.p.). **K:** CD11b+Ly6Gneg macrophage and monocyte plots from the peritoneal cavity of either saline, AC, or AC+AK-treated C57BL/6 mice (AK: Anakinra 30mg/kg i.p.). Data shown represent three or more experiments and are expressed as mean±SEM; Student t-test was used for analysis; P values are indicated in the graphs.

### NLRP3 but not IL-1 signaling is involved in AC-induced changes to the macrophage/monocyte populations in the peritoneal cavity *in vivo*

Having observed a differential response to alum *in vitro*, we next decided to assess the role of NLRP3 in the effects of AC *in vivo*. We performed the conditioning protocol on both C57BL/6 and NLRP3-deficient mice, followed by phenotyping the different populations of macrophages and monocytes. We found that in alum-treated NLRP3^-/-^ mice, pro-caspase-1 did not disappear, suggesting that NLRP3 is required for caspase-1 activation by alum *in vivo* (**Fig. 7C**). Interestingly, AC-mediated MDR happened in lesser extent in the absence of NLRP3 when compared to WT type mice, as observed by a higher number of remaining LPMs in the peritoneal cavity, both in terms of percentage and total cell count (**Fig. 7G**). This suggests MDR is partially dependent on NLRP3 activation. Additionally, we observed that NLRP3 is involved in the migration of SPMs, monocytes, and neutrophils to the peritoneal cavity (**Fig. 7F-I**), indicating that NLRP3 is essential for the inflammatory response triggered by the conditioning protocol.

### AC increases peritoneal macrophages and monocytes susceptibility to *E. coli* infection but boosts inflammatory neutrophil recruitment and bacterial killing *in vitro*

As mentioned, we found that AC induces a major change and long-term change in the immune cell landscape of the peritoneal cavity (add figs). Previous work demonstrates that, upon resolution of inflammation, the newly repopulated peritoneal cavity macrophages present higher bacterial killing (38). To then assess the functional impact of macrophages elicited by AC, we isolated the populations described above and infected them with GFP-expressing *E. coli* at a multiplicity of infection (MOI) of 5 for 24 hours *in vitro*. We found that steady-state LPMs and SPMs were more effective at killing *E. coli* compared to AC-induced SPMs and monocytes (**Fig. 8A**). This diminished bacterial clearance was further confirmed by reduced NO production in AC-induced populations (**Fig. 8B**). Interestingly, this immunosuppressive effect was also observed in LPMs that formed aggregates and those that migrated to the omentum, as evidenced by both higher CFU counts and lower NO production. These results suggest that AC affects all macrophage and monocyte populations within the peritoneal cavity.

**Figure 8:**
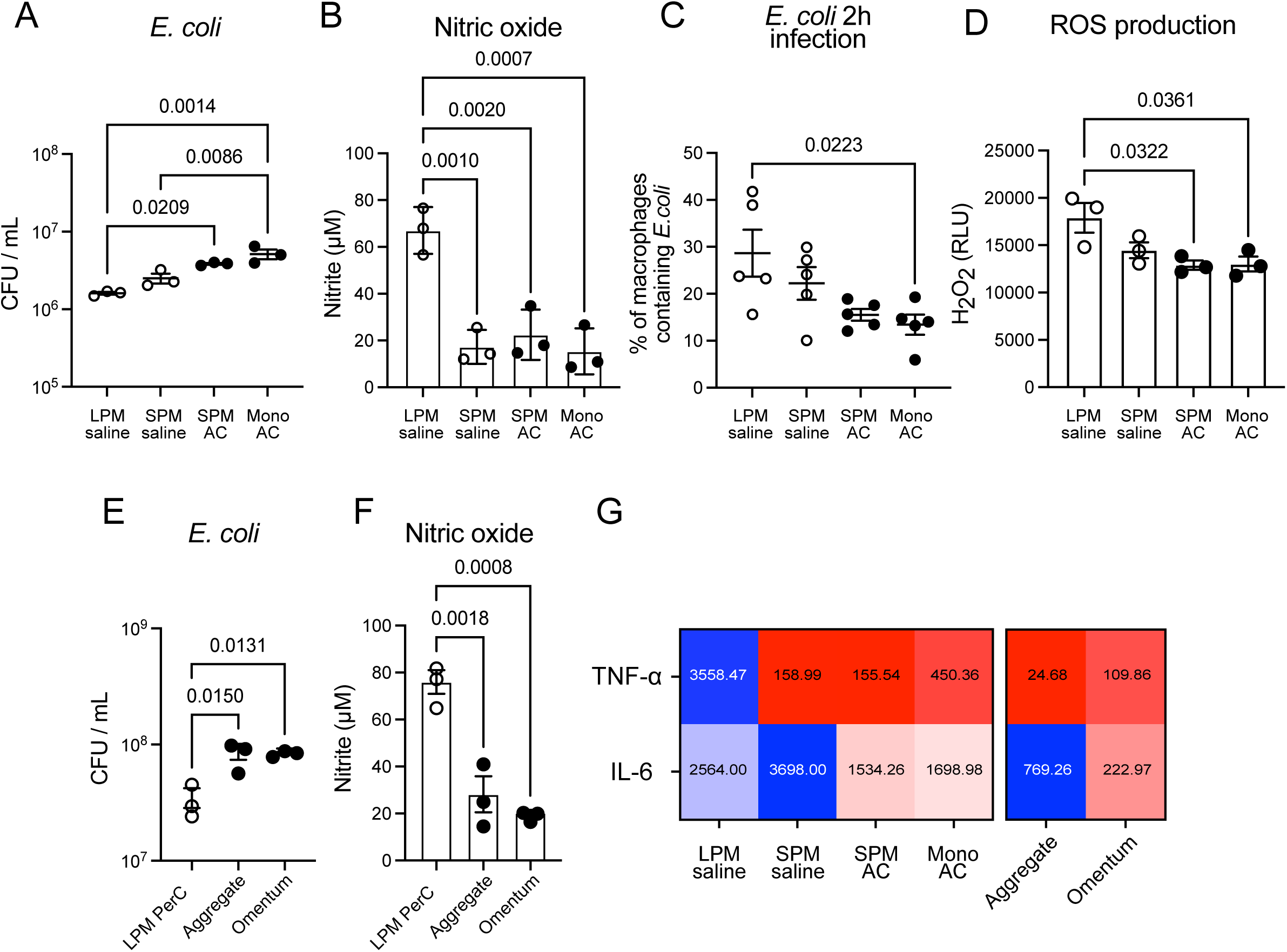
AC renders peritoneal macrophages and monocytes more susceptible to *E. coli* infection *in vitro*. **A:** *E. coli* CFU counts from sorted LPMs, SPMs and monocytes from either saline or AC-treated C57BL/6 mice infected for 24h in an MOI of 5. **B:** NO production from sorted LPMs, SPMs and monocytes from either saline or AC-treated C57BL/6 mice infected for 24h in an MOI of 5. **C:** *E. coli* CFU counts from LPMs sorted from the peritoneal cavity, aggregates or the omentum of either saline or AC-treated C57BL/6 mice infected for 24h in an MOI of 5. **D:** NO production from LPMs sorted from the peritoneal cavity, aggregates or the omentum of either saline or AC-treated C57BL/6 mice infected for 24h in an MOI of 5. **E:** ROS production from sorted LPMs, SPMs and monocytes from either saline or AC-treated C57BL/6 mice infected for 24h in an MOI of 5. **F:** TNF-α and IL-6 production from sorted LPMs, SPMs and monocytes from the peritoneal cavity, aggregates and omentum of either saline or AC-treated C57BL/6 mice infected for 24h in an MOI of 5. Data shown represent three or more experiments and are expressed as mean±SEM; Student t-test was used for analysis; P values are indicated in the graphs.

The subcutaneous administration of alum has been shown to improve pup survival in neonatal sepsis models in an NLRP3-dependent manner, with myeloid GR1+ cells playing a key role in this protection (39, 40). Based on these results and our observations that AC induces major recruitment of neutrophils (Fig 3F), we aimed to functionally assess the neutrophils that migrated to the peritoneal cavity following AC. While AC reprogrammed macrophage and monocyte populations in the peritoneal cavity to a less inflammatory state, its effect on the recruited neutrophils was distinct. Specifically, AC-induced CD11b+Ly6C/G+ neutrophils demonstrated enhanced phagocytic activity (Fig 9A), and more efficient bacterial clearance (Fig 9B) compared to neutrophils recruited by thioglycolate. Furthermore, during both LPS-induced *in vivo* stimulation and infection with *E. coli*, AC-induced neutrophils exhibited a more inflammatory phenotype, characterized by increased ROS and NO production, as well as elevated TNF-α and IL-6 secretion (**Fig 9C-G**). These findings highlight that AC has distinct and differential effects on various cell populations in the peritoneal cavity.

**Figure 9:**
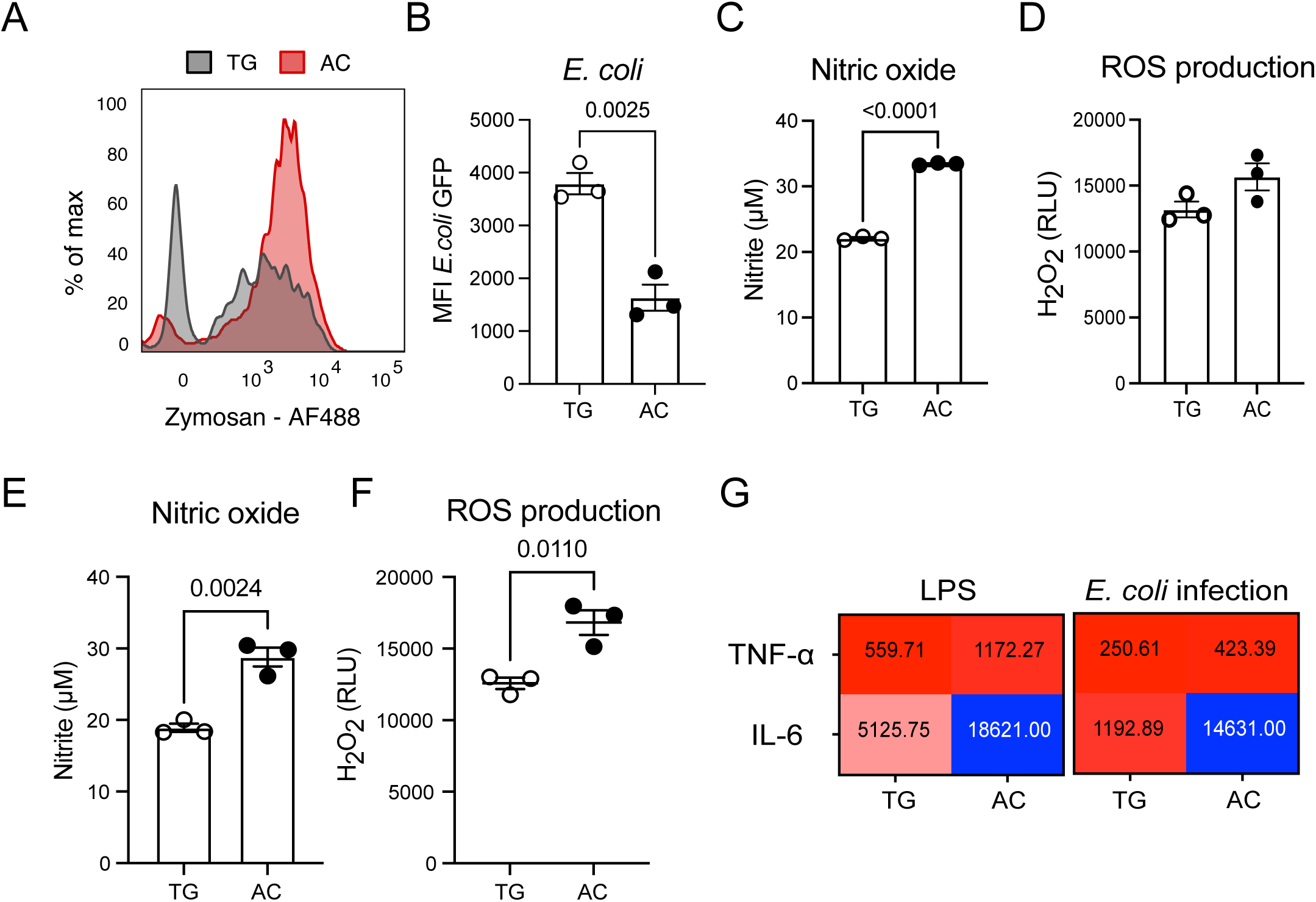
AC boosts inflammatory neutrophil recruitment and bacterial killing *in vitro*. **A:** Histograms of AF488 conjugated Zymosan-A-treated from sorted CD11b^+^Ly6G^+^ cells from thioglycolate or AC-treated mice. Cells were stained for flow cytometry phenotyping and analyzed for phagocytosis. **B:** GFP MFI in sorted CD11b^+^Ly6G^+^ cells from thioglycolate or AC-treated mice infected with *E. coli* GFP for 24h at an MOI of 5. **C:** NO production by sorted CD11b^+^Ly6G^+^ cells from thioglycolate or AC-treated mice infected with *E. coli* GFP for 24h at an MOI of 5. **D:** ROS production by sorted CD11b^+^Ly6G^+^ cells from thioglycolate or AC-treated mice infected with *E. coli* GFP for 24h at an MOI of 5. **E:** NO production by sorted CD11b^+^Ly6G^+^ cells from thioglycolate or AC-treated mice treated *in vitro* with LPS (200ng/mL) for 24h. **F:** ROS production by sorted CD11b^+^Ly6G^+^ cells from thioglycolate or AC-treated mice treated *in vitro* with LPS (200ng/mL) for 24h. **G:** TNF-α and IL-6 production from sorted CD11b^+^Ly6G^+^ cells from thioglycolate or AC-treated mice infected with *E. coli* GFP for 24h at an MOI of 5 or treated *in vitro* with LPS (200ng/mL) for 24h. Data shown represent three or more experiments and are expressed as mean±SEM; Student t-test was used for analysis; P values are indicated in the graphs.

## Discussion

We observed that AC induces a differential response in two subsets of resident peritoneal macrophages: small peritoneal macrophages (SPMs) and large peritoneal macrophages (LPMs). *In vitro*, SPMs exhibited a greater propensity for inflammasome activation, which is consistent with previous reports linking inflammasome activation to proinflammatory responses (41). This suggests that SPMs may play a key role in the initial immune response to alum. Conversely, LPMs showed a more muted inflammasome activation, hinting at a potential functional distinction between these populations in the context of immune modulation. *In vivo*, alum administration led to the reprogramming of peritoneal macrophages and circulating monocytes towards a less inflammatory phenotype. This reprogramming is likely a pivotal step in the establishment of an immunosuppressive environment, which could explain the delay in allogeneic graft rejection observed in previous studies (19, 20). Our results provide new insight into how AC influences immune cells at the site of alum exposure, particularly in the peritoneal cavity, and underscore the importance of macrophage phenotypic shifts in regulating immune responses.

The immunosuppressive effects of AC were not limited to macrophages alone. Our data show that the reprogrammed macrophages were associated with increased susceptibility to bacterial infection, suggesting that the conditioning process dampens the immune system’s ability to mount effective responses against pathogens. This finding is consistent with the well-established trade-off between immune tolerance and immune defense, where an immunosuppressive environment may reduce inflammatory responses to allogeneic tissues but also impair defense against infections (42, 43).

Despite these findings, while macrophages became less inflammatory, AC simultaneously recruited neutrophils with higher killing ability. This suggests that AC may fine-tune the immune response, balancing suppression of allogeneic rejection and pathogen defense by enhancing neutrophil functionality (39, 40) while suppressing the inflammatory activity of macrophages. This dual effect highlights the complexity of immune reprogramming and suggests that AC may be a promising therapeutic approach to modulate immunity in conditions such as transplant rejection and autoimmune diseases, where immune tolerance to self or transplanted tissues is required, yet pathogen defense must be preserved.

While our *in vitro* findings are important, they may not fully represent the *in vivo* processes that occur during trauma or sterile inflammation. To better understand the role of NLRP3 specifically in myeloid cells *in vivo*, further experiments using conditional knockout mice are necessary. These studies could help determine whether this signaling pathway is involved in sepsis models and if the immunosuppressive phenotype induced in macrophages could protect in allogeneic transplantation models. Additionally, while clodronate liposomes are a valuable tool for studying the origins of both SPM and LPM, they may also deplete myeloid cells that arise during emergency myelopoiesis, such as MDSCs. Given that our work shows AC induces the expansion of functional MDSCs, which play a role in shaping the adaptive immune response and delaying allograft rejection, depleting macrophages could potentially affect MDSC expansion as well.

In conclusion, our findings add new dimensions to our understanding of AC and its ability to reprogram the immune system. The differential response of SPMs and LPMs to alum exposure and the subsequent effects on both innate and adaptive immunity underscore the complexity of the immune modulation process. While the therapeutic potential of AC in managing autoimmune diseases and transplant rejection is promising, further research is needed to delineate the specific mechanisms underlying the reprogramming of immune cells and to explore strategies that could mitigate the risk of increased infection susceptibility. Future work could also investigate the long-term effects of AC on immune homeostasis, as well as its interactions with other immune-modulating therapies. The development of targeted interventions that selectively modulate immune cell subsets without compromising host defense will be crucial for the successful application of AC in clinical settings.

## Acknowledgements

This work was supported by the American Pediatric Surgical Association Jay Grosfeld Scholar Grant, the Society of University Surgeons Junior Faculty Award, the Hardy Hendren Faculty Development Fund at Boston Children’s Hospital, the Junior Translational Investigator Service Award from the Translational Research Program at Boston Children’s Hospital.

Efforts of VF were funded by the Assistant Secretary of Defense for Health Affairs endorsed by the Department of Defense through the Peer Reviewed Medical Research Program under award #HT9425-23-1-0040 as well as by the National Institutes of Health (NIH) through the NIH HEAL Initiative (https://heal.nih.gov/) under award #K99HD115239. Opinions, interpretations, conclusions, and recommendations contained herein are those of the authors and are not necessarily endorsed by the Department of Defense or NIH. MDVdS thanks Coordination for the Improvement of Higher Education Personnel (CA PES) for the 12-month scholarship to develop his Split Fellowship (Doutorado Sanduíche PDSE).

## Supplemental Figure Legends

**Supplemental Figure 1:** 3×10^5^ peritoneal cells from C57BL/6 mice were plated on 96-well plates overnight. After adhesion, non-adherent cells were washed out with cold PBS, and cells were primed with LPS (50ng/mL) for 2h and subsequently stimulated with alum Imject at 250, 500 or 1000μg/mL for 6h. Cells were assessed for viability with EtBr^+^ staining.

**Supplemental Figure 2. A:** gating strategy used for analyzing macrophage populations after collection from the peritoneal cavity. **B:** Sorted SPMs and LPMs plated at 2×10^5^ cells/well, were primed with LPS (50ng/mL) for 2h and subsequently stimulated with alum Imject at 500μg/mL for 6h. Cells were assessed for viability with EtBr^+^ staining.

**Supplemental Figure 3.** Analysis of macrophages and monocyte populations in the peritoneal cavity, omentum and aggregates of either saline or AC-treated mice at 1 day, 7 days and 14 days after the last injection.

