## Supplementary figures and images for "Adjuvant conditioning enhances neutrophil function while inducing a suppressive peritoneal macrophage phenotype"

### Supplemental Figures 1-3

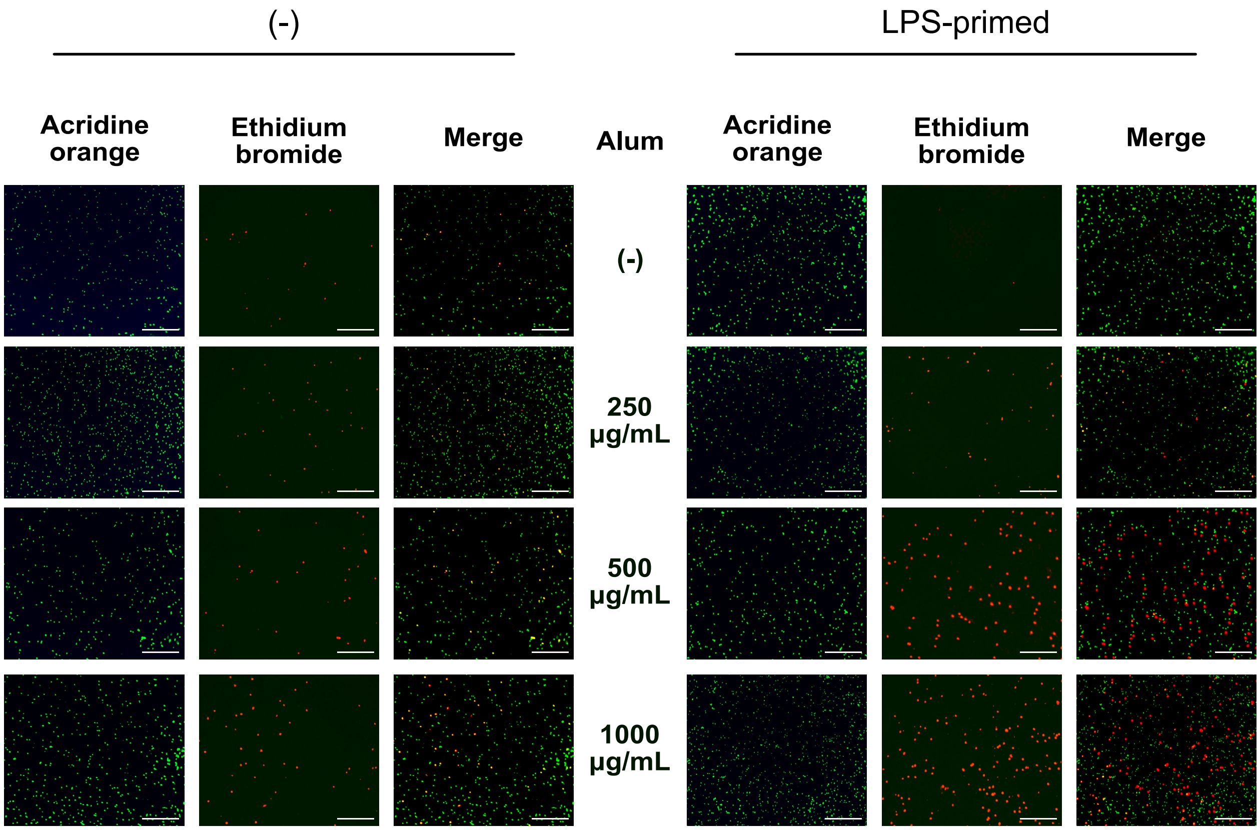

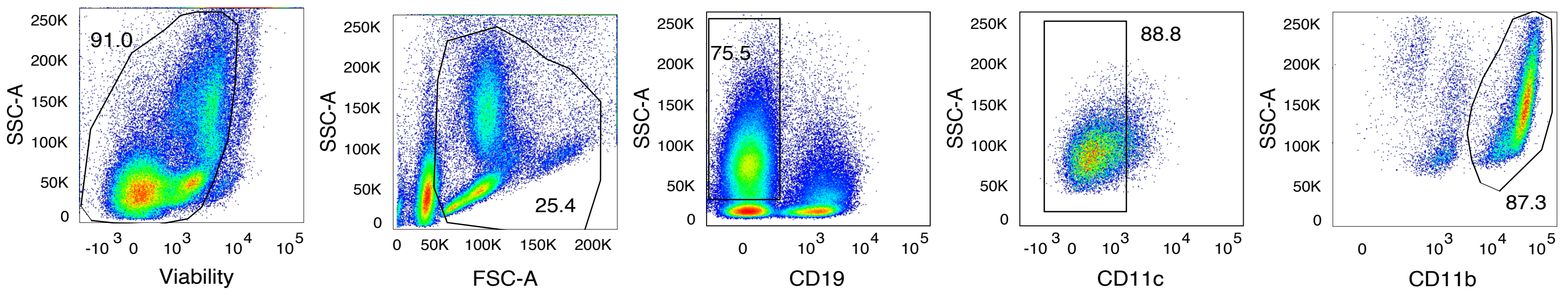

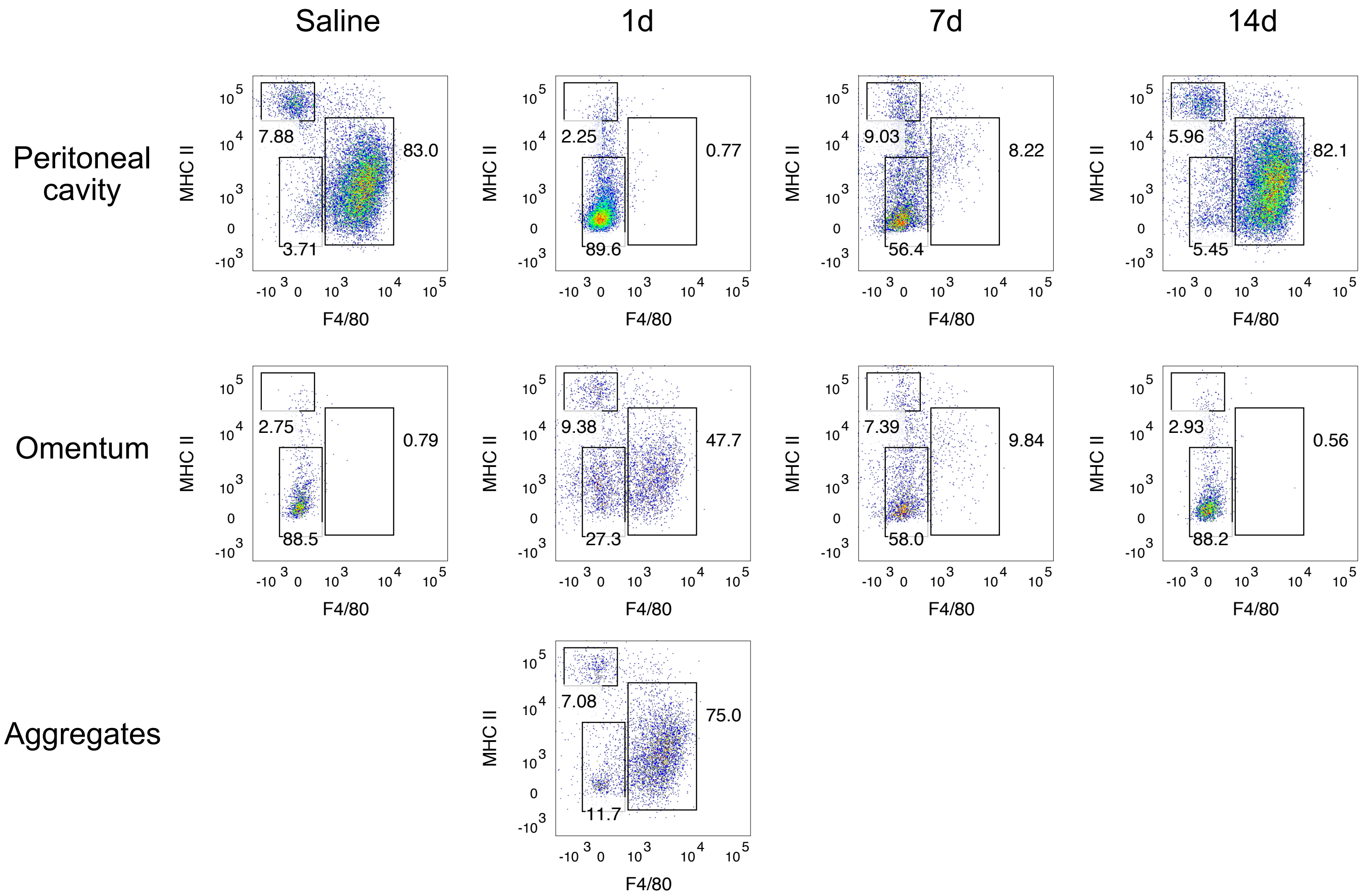
